# Break-induced replication is the primary recombination pathway in plant somatic hybrid mitochondria: a model for mt-HGT

**DOI:** 10.1101/2023.01.07.523103

**Authors:** Carolina L. Gandini, Laura E. Garcia, Cinthia C. Abbona, Luis F. Ceriotti, Sergei Kushnir, Danny Geelen, M. Virginia Sanchez-Puerta

## Abstract

Somatic hybrids between distant species offer a remarkable model to study genomic recombination events after mitochondrial fusion. Recently, our lab described highly chimeric mitogenomes in two somatic hybrids between the Solanaceae *Nicotiana tabacum* and *Hyoscyamus niger* resulting from interparental homologous recombination. To better examine the recombination map in somatic hybrid mitochondria, we developed a more sensitive bioinformatic strategy to detect recombination activity based on high-throughput sequencing without assembling the hybrid mitogenome. We generated a new intergeneric somatic hybrid and re-analyzed the two Solanaceae somatic hybrids. We inferred 213 homologous recombination events across repeats of 2.1 kb on average. Most of them (∼80%) were asymmetrical, consistent with the break-induced replication (BIR) pathway. Only rare (2.74%) non-homologous events were detected. Interestingly, independent events frequently occurred in the same regions within and across somatic hybrids, suggesting the existence of recombination hotspots in plant mitogenomes. BIR is the main pathway of interparental recombination in somatic hybrid mitochondria. Likewise, under the fusion compatibility model of mitochondrial horizontal transfer, foreign mitochondria fuse with those in the recipient cell and their genomes likely recombine via BIR, resulting in the integration and/or loss of mitochondrial DNA. Findings of this study are also relevant to mitogenome editing assays.

**Highlight:** We show that the chimeric mitochondrial genomes of somatic hybrids result from one of the three described homologous recombination pathways (BIR), mimicking the fusion compatibility model for plant HGT.

## Introduction

In all domains of life, DNA is vulnerable to damage by endogenous insults or environmental agents that harm the replication processes and the fidelity of the genetic transmission (Lindahl *et al*., 1993). The principal cytotoxic lesions are DNA double-strand breaks (DSBs). The repair of DSBs can occur in two ways: non-homologous pathways involving the ligation of the broken DNA ends with none or little similarity, or homologous recombination (HR), requiring the availability of homologous regions as template for the repair (Daley *et al*., 2005; Mehta and Haber, 2014). HR can take place through alternative pathways depending on the structure of DNA ends and the intrinsic activity of the DSB repair machineries, yielding different products of recombination. When both ends of the DSB share homology with a template sequence, either synthesis-dependent strand annealing (SDSA) or double Holliday junction (dHJ) pathways can operate (Mehta and Haber, 2014). During SDSA the invading strand is displaced after DNA synthesis and anneals with the second end leading to exclusively non-crossover (non-CO) products and a non-reciprocal transfer of genetic information (Mehta and Haber, 2014). Conversely, the dHJ pathway yields to crossover (CO) or non-CO products depending on the resolution/dissolution mechanism (Resnick, 1976; Szostak *et al*., 1983; Wu *et al*., 2003). When only one end of the DSB shares homology with the donor sequence or a replication fork is stalled, the break-induced replication (BIR) pathway becomes necessary leading to non-reciprocal recombination (Malkova *et al*., 1996; Sakofsky and Malkova, 2017).

Accumulating evidence supports the existence of mitochondrial recombination in plants; however, only a few proteins involved in mitogenome recombination have been characterized using *Arabidopsis* and *Physcomitrella* nuclear mutants (Chen, 2013; Odahara *et al*., 2015, 2017, 2021; Gualberto and Newton, 2017; Chevigny *et al*., 2020). Mutants of nuclear factors with a role in recombination surveillance, such as MSH1 and OSB1, often show an increase in mitogenome rearrangements due to asymmetrical recombination across repeats of intermediate size (Zaegel *et al*., 2006; Arrieta-Montiel *et al*., 2009; Miller-Messmer *et al*., 2012). In other cases, an accumulation of chimeric molecules that emerge from microhomology-mediated repair was observed in the mitochondria of mutants of factors involved in HR-dependent repair, such as RECA or WHY2 (Cappadocia *et al*., 2010; Davila *et al*., 2011; Odahara *et al*., 2021). Non-homologous repair activity is only exceptionally reported in wild-type plants (Zou *et al*., 2022) and can lead to structural changes among different species or varieties (Davila *et al*., 2011; Cole *et al*., 2018; Zou *et al*., 2022) or to gene chimeras resulting in cytoplasmic male sterility phenotypes (Schnable, 1998). In general, most recombination events in plant mitochondria go unseen because DSBs are repaired by HR using as template an identical copy of the mitogenome. However, non-allelic HR across large repeats (> 1 kb) is frequent and leads to alternative forms of the mitogenomes observed in Southern blots (Stern & Palmer, 1984) or using long read sequencing technologies (Fischer *et al*., 2022; Zou *et al*., 2022). In contrast, intermediate and small repeats rarely recombine as ectopic recombination across these repeats is suppressed by nuclear factors (Davila *et al*., 2011). These results imply that both homologous and, more rarely, non-homologous pathways exist in plant mitochondria (Stern and Palmer, 1984; Kanazawa *et al*., 1998; Scotti *et al*., 2004; Zaegel *et al*., 2006; Allen *et al*., 2007; Shedge *et al*., 2007; Odahara, 2020). Much work is still needed to clarify the mechanisms of DNA recombination, repair, and replication (RRR) and their relative importance in wild type plant mitochondria.

An excellent model for studying recombination across repeats in plant mitochondrial genomes are somatic hybrids, which result from the protoplast fusion of two different species. Their particular nature allows the study of mitochondrial recombination events given that parental mitochondria undergo fusion (Oldenburg and Bendich, 1996, 2015; Arimura *et al*., 2004) leading to a plethora of homologous regions which act as non-identical repeats of variable sizes and sequences within the mitochondria of the somatic hybrid. Multiple intergenomic recombination events across these repeats (non-allelic HR) can be easily identified by DNA sequencing. Even though earlier studies have described the chimeric nature of the mitogenome of somatic hybrids (Belliard *et al*., 1979; Vedel *et al*., 1986; Zubko *et al*., 1996), recombination events have only been thoroughly studied in three cytoplasmic hybrids by assembling the mtDNA and comparing it with those of their parents (Sanchez-Puerta *et al*., 2015; Garcia *et al*., 2019; Vasupalli *et al*., 2021). However, assembling plant mitogenomes is challenging due to the presence of repeated sequences and only a subset of the co-existing re-arranged molecules are represented in a given assembly (Kmiec *et al*., 2006; Fischer *et al*., 2022), underestimating the real number of recombined molecules in the cybrids. To overcome these difficulties, we developed a bioinformatic pipeline that infers recombination events by analyzing mapping patterns and genotype swaps within paired-end reads without assembling the mitogenome of the hybrid. Furthermore, the DNA repair pathway responsible for the HR events may be inferred from the precisely characterized products of recombination. To examine a greater sample of recombination events, we produced a somatic hybrid by chemical protoplast fusion between *N. tabacum* and another Solanaceae, *Physochlaina orientalis* and evaluated the mitochondrial recombination map of leaf and callus tissue. In addition, we re-analyzed two previously studied somatic hybrids *N. tabacum* + *Hyoscyamus niger* in a search of recombination hotspots.

## Materials and Methods

### Somatic hybrid production, selection, and sequencing

Somatic hybrid production was achieved by protoplast fusion between an albino *N. tabacum* CAT-9 line and *P. orientalis* (Fig. 1). Tobacco CAT-9 is a transplastomic line, which carries a codon replacement in the plastid gene *atpA* (Schmitz-Linneweber *et al*., 2005). Seeds from *P. orientalis* (NBG944750045) were obtained from the Nijmegen Botanical Garden (The Netherlands). Plants of both species were propagated *in vitro* on basal Murashige and Skoog (MS) medium at 25°C and 8/16 h dark/light photoperiod. *N. tabacum* and *P. orientalis* protoplasts were isolated from leaves of aseptically grown plants using an enzymatic solution; the mixed protoplast suspension was chemically fused using PEG fusion solution (Methods S1). After fusion, the protoplasts were cultivated in K3 medium with decreasing concentrations of D-Mannitol and with hormones (Methods S1). After two months, calli were plated on the surface of an MSR medium solidified with 0.7% agar until green colonies appeared, regenerating shoots were transferred to MS medium. For Southern blot assays, DNA was extracted from the parental lines (*N. tabacum* and *P. orientalis*) and from green calli, digested with *SacI* (Promega) and separated by electrophoresis in 0.8% agarose (Methods S1). Southern blot hybridization was performed with probes of the mitochondrial genes *atp1, cob*, and *cox1*. Calli in which mitochondrial recombination was confirmed (Fig. S1) were cultured in MS medium for plant regeneration. Total DNA extracted from calli and leaves of a selected somatic hybrid line named ntpoA01 was subjected to high-throughput sequencing using the Illumina HiSeq 2500 platform at the Beijing Genomics Institute. A total of 6.6 Gb data containing ∼26.5M clean 2×125 bp paired-end (PE) reads with an average insert size of ∼800 bp were generated. Clean sequence data are available from the NCBI Bioproject ID PRJNA899353. The chloroplast genome of the ntpoA01 somatic hybrid was assembled using GetOrganelle (Jin *et al*., 2020), with standard parameters and *Nicotiana sylvestris* (NCBI accession NC_007500) as seed. The assembled sequence can be found as Supplementary Material (Data S1).

**Figure 1.**
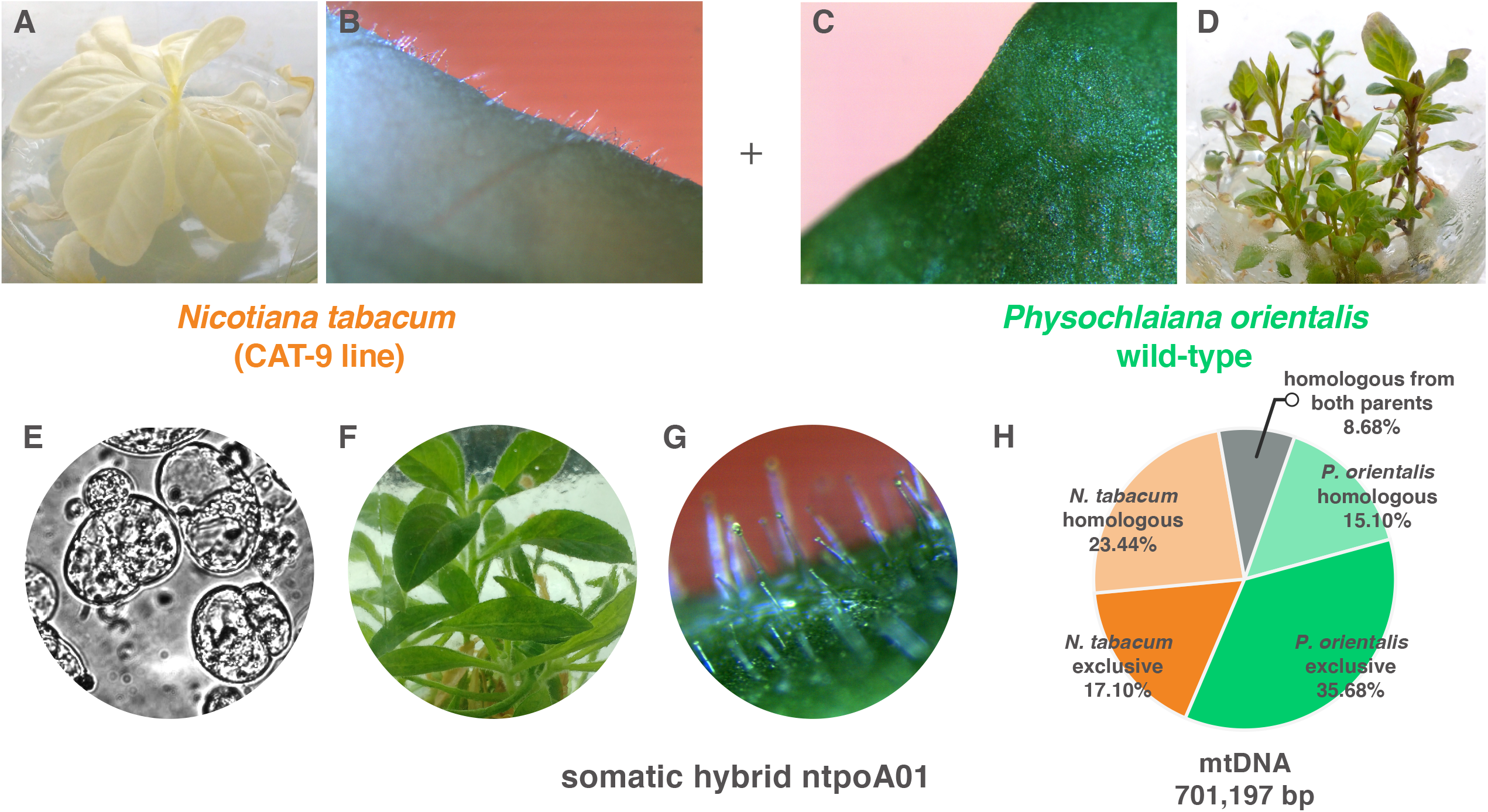
Production of the ntpoA01 somatic hybrid plant. **A,B)** Image of the parental *N. tabacum* albino (CAT-9 line) plant, and close-up of a leaf showing the presence of trichomes. **C)** *P. orientalis* leaves are trichome-less. **D)** Image of the parental *P. orientalis*. **E)** Cell divisions after protoplast fusions between *N. tabacum* and *P. orientalis*. **F,G)** Image of the ntpoA01 regenerated plant and its leaves are hairy as tobacco. **H)** Pie chart depicting the parental contribution to the mitochondria of ntpoA01.

### Comparison of parental mitogenomes

Homologous regions between the parental mitogenomes (NCBI NC_006581.1 and NC_044153.1) were identified using BLASTn with default presets. Hits greater than 25 bp in length and 80% of sequence identity were further analyzed. Blast hits located in the genomes at a distance shorter than the library inner size (550 bp) were joined. In addition, identical homologous regions between the two parental mitogenomes were identified using VMATCH (Kurtz, 2003). Parental-exclusive regions were calculated by subtracting homologous regions from the corresponding mitogenome. A single copy of repeats larger than 1 kb within each parental mitogenome was considered for comparative and recombination analyses.

### Detection of recombination events

To examine the chimeric molecules in the somatic hybrid ntpoA01 mitochondria, we developed a novel assembly-free bioinformatic strategy that detects recombination events from genomic data. The strategy involves several steps, described below, and detailed in Fig. S2:

1. Read mapping: a total of 943,990 paired-end (PE) reads mapped to the tobacco and *P. orientalis* parental mitogenomes, using Bowtie2 (Langmead and Salzberg, 2012) end-to-end alignment with default presets. Only best alignments were considered.
2. Read filtering: Three datasets were created by differential filtering the Bowtie2 alignment:
  a. Concordant PE-reads that aligned perfectly: only reads with their read pair properly mapped (*i*.*e*., aligned to the same genome and at a distance equal to the insert size +/- 400 bp)) and exhibiting a 100% identity with the reference were considered. The *N. tabacum* and *P. orientalis* alignments yielded 107,365 and 202,264 PE-reads, respectively.
  b. Discordant PE-reads perfectly mapped: only reads with their read pair mapped to the other parental mitogenome and with a 100% identity to the reference were considered. A total of 99,472 PE-reads were perfectly aligned between tobacco and *P. orientalis* parental mitogenomes.
  c. Informative PE-reads: this is a subset of those described in 2b; we eliminated those read pairs in which at least one of the reads aligns to an identical homologous region between the parental mitogenomes. This yielded a total of 4,292 PE-reads (only 4.31% of the discordant PE-reads).
3. Inference of parental contribution: concordant and discordant PE-reads perfectly mapped to either parental mitogenome were used to infer the minimal parental contribution to the ntpoA01 mitogenome (parental regions that were kept in ntpoA01). A parental region was inferred to be present in the somatic hybrid if it spans a minimum length of 1 kb with a read coverage equal to or greater than three.
4. Evidence of recombination: recombination events in the hybrid were inferred from the informative PE-reads by detecting genotype switches between parental mitogenome. A breakpoint region was defined as the shortest fully identical genomic region between the two parents flanked by single nucleotide polymorphisms (SNPs). Evidence of homologous recombination results from a minimum of three read pairs in which:
  a. One or both of the reads of a read pair map to a non-identical repeat region shared by the parental lines greater than the inner size (*i*.*e*., > ∼550 bp) or,
  b. Both reads maps at opposite ends of a homologous region shorter than the inner size (*i*.*e*., < ∼550 bp).
  In contrast, evidence for non-homologous events result from informative PE-reads that mapped to regions in each parental mitogenome that are not linked by homologous tracts. These events were inferred when at least three informative PE-reads were found in the same breakpoint region.

With the assembly-free methodology, the somatic hybrids between *N. tabacum* and *Hyoscyamus niger* (nthn) (NCBI accessions SRR1509957.2 and SRR8445683) were re-analyzed (Table S1). Scripts used in this analysis can be found in https://github.com/cgandini/somatic_hybrids.

Plots and figures were generated using R v.3.5.0, circos v.0.69-8 and/or Adobe Illustrator 2019.

### Statistical analyses of recombining repeats

A Student’s t-Test to asses’ differences in length and sequence similarity (identity) between recombining and non-recombining homologous regions was performed in R using the *t*.*test* function included in the *stats* package. Statistical association between recombinational breakpoints occurring in the nthn somatic hybrids was conducted using the *RegioneR* R package v.1.29.1 (Gel *et al*., 2015), which implements a permutation test framework specifically designed to test correlation between two sets of genomic regions. The tests were performed using the *overlapPermTest* function with the parameters: ntimes=10,000 and alternative = “greater”. Both parental mitogenomes were masked so permutation was allowed only over homologous regions between parental mitochondria satisfying a minimum length of 99 bp and minimum identity of 87.50% (as these are the minimum values for recombining repeats in this study).

## Results

### Tobacco and *Physochlaina orientalis* somatic hybrid production and regeneration

The ntpoA01 line selected for this study was produced by protoplast fusion between two Solanaceae species. First, somatic hybrids were produced by chemical protoplast fusion between an albino *N. tabacum* CAT-9 line and a wild-type individual of *P. orientalis*, which belong to two distant tribes of the family Solanaceae (Olmstead *et al*., 2008) and are sexually incompatible species. The combination of two selectable markers, the green pigmentation from one parent and the higher regeneration rate from the other, allowed the identification of the somatic hybrids. Hybrid lines were evaluated for mitochondrial chimerism using Southern blots (Methods S1, Fig. S1). The somatic hybrid line A01 with a hybrid mitogenome was selected. The regenerated plant is phenotypically similar to tobacco in leaf morphology, geometry and trichome presence, suggesting the prevalence of the tobacco nuclear genome (Fig. 1) while it exhibits green pigmentation indicating that the chloroplast is derived from *P. orientalis*. A comparative analysis of the ntpoA01 chloroplast genome showed that it was identical to that of *P. orientalis* (Data S1).

### Parental mitogenome comparisons and their contribution to the ntpoA01 mitogenome

The mitochondrial genomes of both parental lines used in protoplast fusions are available for genomic comparisons. The *P. orientalis* line is the same as the one used for mitogenome sequencing (Gandini *et al*., 2019) and the mtDNA of *N. tabacum* CAT9 (unpublished results) is identical to the published tobacco mitogenome (Sugiyama *et al*., 2005). Despite an almost identical gene content (Table S2), the mitogenome of *P. orientalis* is longer than that of *N. tabacum* (684,857 bp versus 430,597 bp) and contains more duplicated regions. Thus, to avoid overestimations in all comparisons and calculations, repeats greater than 1 kb were considered only once, resulting in parental genome lengths of 396,065 bp and 575,183 bp for *N. tabacum* and *P. orientalis*, respectively. Two categories of sequences can be established when comparing the parental genomes: *(i)* those shared by the two parents, *i*.*e*., homologous regions, which become repeats within the somatic hybrid mitochondria; and *(ii)* those exclusive to each parent, *i*.*e*., exclusive regions (Table S3). Briefly, about 267 and 274 kb of DNA is shared by both mitogenomes (representing ∼67% of *N. tabacum* and ∼48% of *P. orientalis*, respectively) including a total of 1,106 homologous regions (840 and 943 considering only once those that fall within intraparental repeats in *N. tabacum* and *P. orientalis*, respectively) ranging from 26 bp to ∼7,558 bp in length and from 80.56% to 100% in sequence identity (Table S3, Table S4). Among the homologous regions only 132 are identical between the parental genomes with a maximum length of 339 bp. On the other hand, a total of ∼129 kb and ∼301 kb are exclusive regions in *N. tabacum* and in *P. orientalis* mitogenomes, respectively.

By analyzing the ntpoA01 concordant PE-reads that align perfectly to each parental mitogenome we examined how many of the exclusive and homologous regions from each parental line are present in the somatic hybrid mitochondria (Fig. 1H). Almost all exclusive sequences from each parent (∼92% of *N. tabacum* exclusive regions and ∼83% of those exclusive to *P. orientalis*) are retained in ntpoA01. *N. tabacum* contributed with more homologous regions than *P. orientalis*, 194,792 bp (∼73%) versus 136,312 bp (∼49%) (Table S3). In summary, the somatic hybrid mitogenome has a minimum length of 701,197 bp (*i*.*e*., ∼28% less than the sum of both parental mitogenomes), lost ∼14% of the exclusive regions, and conserved ∼60% of the homologous regions, which derived mainly from only one of the parents. Regarding the gene content, the ntpoA01 mitogenome retained at least one copy of all genes present in the parental genomes (48 from tobacco and 31 from *P. orientalis*). A total of four protein-coding genes and eight tRNA genes are duplicated as both parental alleles co-exist (Table S2). Of particular note, the intron-containing gene *cox1* was inherited from *P. orientalis*, while the intronless *cox1* from tobacco was lost. In addition, three of the protein-coding genes (*atp1, rps3, rps4*) and one rRNA gene (*rrnL*) are chimeric as a result of recombination between the parental sequences.

### The ntpoA01 mitogenome underwent multiple recombination events, most of which were asymmetrical

To achieve a comprehensive characterization of the interparental HR events in the somatic hybrid ntpoA01, we developed a bioinformatic strategy that identifies recombination events by analyzing parental mitogenome sequence switches in PE-reads. Through the mapping of the ntpoA01 informative PE-reads over the parental mitogenomes (Fig. S2) we were able to infer HR events if a switch fell within a homologous region; otherwise, we considered it a non-homologous event. We detected a total of 115 HR and two non-homologous events (Table S5). Noticeably, HR events occurred only across 54 of the 1,106 homologous regions shared by the parents. That is, multiple independent events were identified at different breakpoints within 31 of the recombining homologous regions (57.41%). The recombining repeats were 99 to 7,523 bp in length and exhibited an average identity of 97.36%.

To shed light into the HR mechanism involved in each event, we analyzed the ntpoA01 concordant and informative PE-reads mapped to each parental homologous region and its flanking areas (See Fig. S3 and Methods S2). This allowed us to characterize the products of recombination pointing to the responsible molecular pathways as illustrated in Fig. 2 on three examples. The entire list of HR events is in Table S5 and its graphical representation for each recombined region is detailed in Fig. S4. Overall, of the 115 HR events, we observed 16 (13.91%) non-crossover products, 4 reciprocal crossovers (3.48%), and 95 asymmetrical recombination events (82.61%).

**Figure 2.**
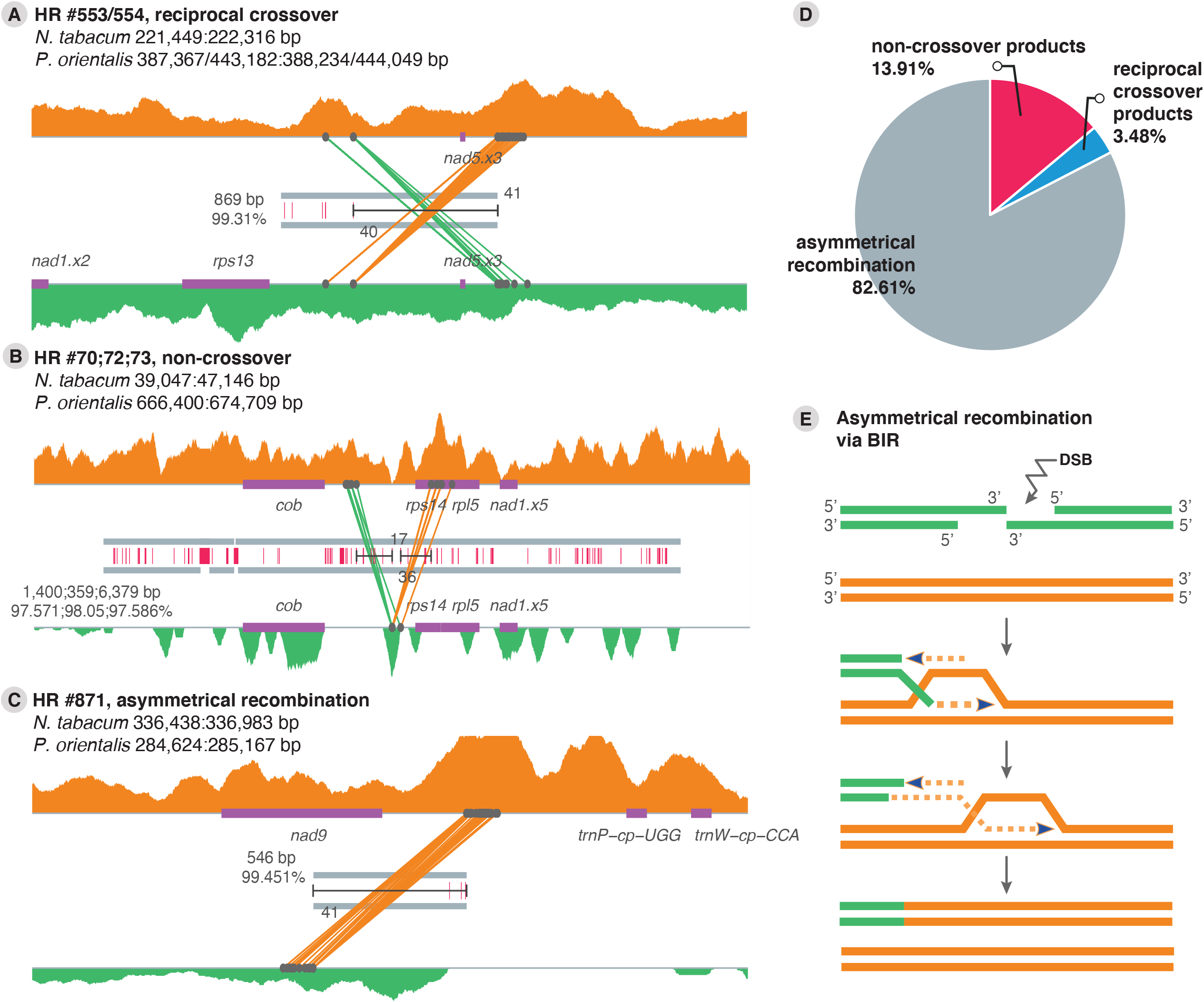
A, B, C) Examples of recombination events in the ntpoA01 mitogenome. The thick gray lines in the middle depict homologous regions (non-identical repeats) between the parental genomes and SNPs between them are denoted with red lines. The thin gray lines represent the *Nicotiana tabacum* (top) and *Physochlaina orientalis* (bottom) mitogenomes with genes shown as violet boxes and including the homologous region and 1,000 bp of its flanking areas. The read depth of parental DNA in the somatic hybrid ntpoA01 determined by concordant PE-reads is shown with orange and green area plots for *N. tabacum* and *P. orientalis*, respectively. Breakpoint regions are shown with a horizontal black line and the number of informative PE-reads are indicated and plotted as diagonal lines that connect both genomes. The length and identity of each homologous region are shown on the left. The identification number of each recombination, the type of recombination products, and the coordinates of the homologous region for each parental mitogenome are detailed at the top of each plot. **D)** Pie chart illustrating the percentage of each recombination type inferred in the ntpoA01 hybrid genome. **E)** Asymmetrical recombination as a result of the break-induced replication (BIR) pathway.

### Reevaluation of two previously studied somatic hybrids using the assembly-free strategy

To evaluate the developed assembly-free methodology and test its sensitivity in comparison with the assembly-based strategy, we reanalyzed the two somatic hybrid mitochondria previously reported (Sanchez-Puerta *et al*., 2015; Garcia *et al*., 2019). We mapped and examined the aligned PE-reads of the two cybrid lines produced from the Solanaceae tobacco and *Hyoscyamus niger* (nthnFH4 and nthnMv-1-1g, respectively) (Table S1). The estimated minimum mitogenome length is 559,846 bp for nthnFH4 and 607,628 bp for nthnMv-1-1g, *i*.*e*., 65% and 70% of the sum of both parental mitogenomes (Table S3). Both nthn cybrids conserved ∼73-97% of the exclusive regions of each parent and have eliminated ∼50% of the homologous regions, most of them from *H. niger* (Table S3). These values are similar to those previously reported (Sanchez-Puerta *et al*., 2015; Garcia *et al*., 2019).

Regarding the recombination analysis, a total of 704 homologous regions were identified between both parental mitogenomes (604 and 639 considering once those that fall within intraparental large repeats in *N. tabacum* and *H. niger*, respectively) (Table S3, Table S6). For nthnFH4, the assembly-free strategy detected 53 HR events in 42 regions and no non-homologous events (Fig. 3, Table S7 and, graphical interpretation in Fig. S5). The homologous regions that recombined ranged from 100 to 9,036 bp in length and from 89.42% to 99.48% in identity. Approximately 31% of the recombining regions were involved in more than one recombination event. Seven of the 53 recombination (13.21%) events led to non-crossover products and 46 (86.79%) to asymmetrical events. No evidence of products of reciprocal crossovers was observed. For the nthnMv-1-1g, a total of 44 HR events in 31 regions were inferred. The homologous regions that recombined ranged from 207 to 9,036 bp in length and from 89.42% to 99.48% in identity, with ∼35% of the recombining regions presenting more than one event (Fig. 3, Table S8 and, graphical interpretation in Fig. S6). A total of three reciprocal crossovers, ten non-crossovers and 31 asymmetrical recombination events were inferred. Also, four non-homologous events were identified.

**Figure 3.**
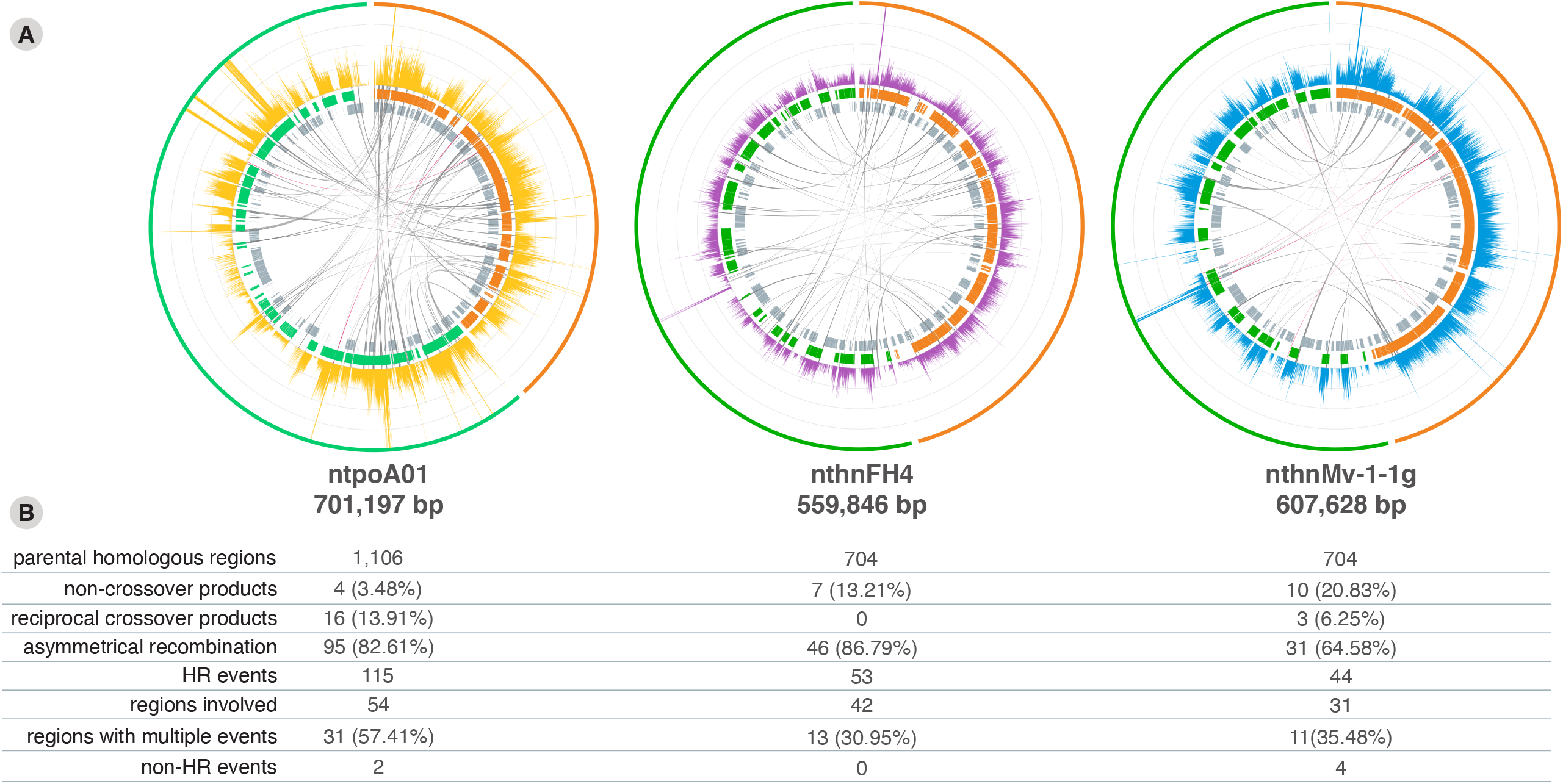
Summary of recombination events in the three somatic hybrid mitogenomes. **A)** From the outer ring to the inner ring: the reference mitogenomes (orange lines for tobacco, light green line for *Physochlaina orientalis*, and dark green lines for *Hyoscyamus niger*); depth of coverage by concordant PE-reads of the hybrids over each parental mitogenome; thick orange or green lines depict parental regions conserved in each hybrid mitogenome; parental homologous regions; homologous recombination and non-homologous events are shown in gray and red links, respectively. The estimated total length of the resulting hybrid genome is showed beneath each plot. **B)** Summary table of recombination events in the somatic hybrids. (HR) Homologous recombination.

The major difference between the assembly-free analysis and the one based on the assembled mitogenomes is that the former has higher sensitivity to detect recombination events, while only those rearrangements found in higher stoichiometries are depicted in the mitogenome assemblies. The strategy designed here was able to infer ∼89% (27) more HR events in the nthnFH4 mitochondria and ∼57.14% (17) more events in nthnMv-1-1g (Table S7 and Table S8). Only one and four events previously detected in the nthnFH4 and nthnMv-1-1g cybrids were not detected by this study, respectively. Given our strict mapping and filtering strategy, four of them fell below our minimum of three read-threshold presenting only two PE-reads of evidence and one have no PE-read of evidence.

### Homologous regions that recombine are larger and more similar than those that did not recombine

While a large number of homologous regions (*i*.*e*., repeats) of different lengths and identities were present in the somatic hybrids, only a subset participated in non-allelic HR events. We tested whether there was a bias in length or sequence identity in those recombining repeats compared to all available repeats in the initial hybrid mitochondria. For this, we used an unequal variance *t*-test for each somatic hybrid testing recombining versus non-recombining regions in terms of length and sequence identity (Fig. 4). Given that the shortest recombining region found in this study is 99 bp long (Table S5), we only considered homologous regions greater or equal to that length. However, the same analysis using all homologous regions shows even greater significance of results (Fig. S7). We found that recombining homologous regions (repeats) are significantly larger and exhibit significantly higher identity than those repeats that did not recombine. Recombination events involve non identical repeats with a mean length of 1864 bp, 2209 bp, and 2242 bp, for each of the three somatic hybrids, respectively (Fig. 4A,B).

**Figure 4.**
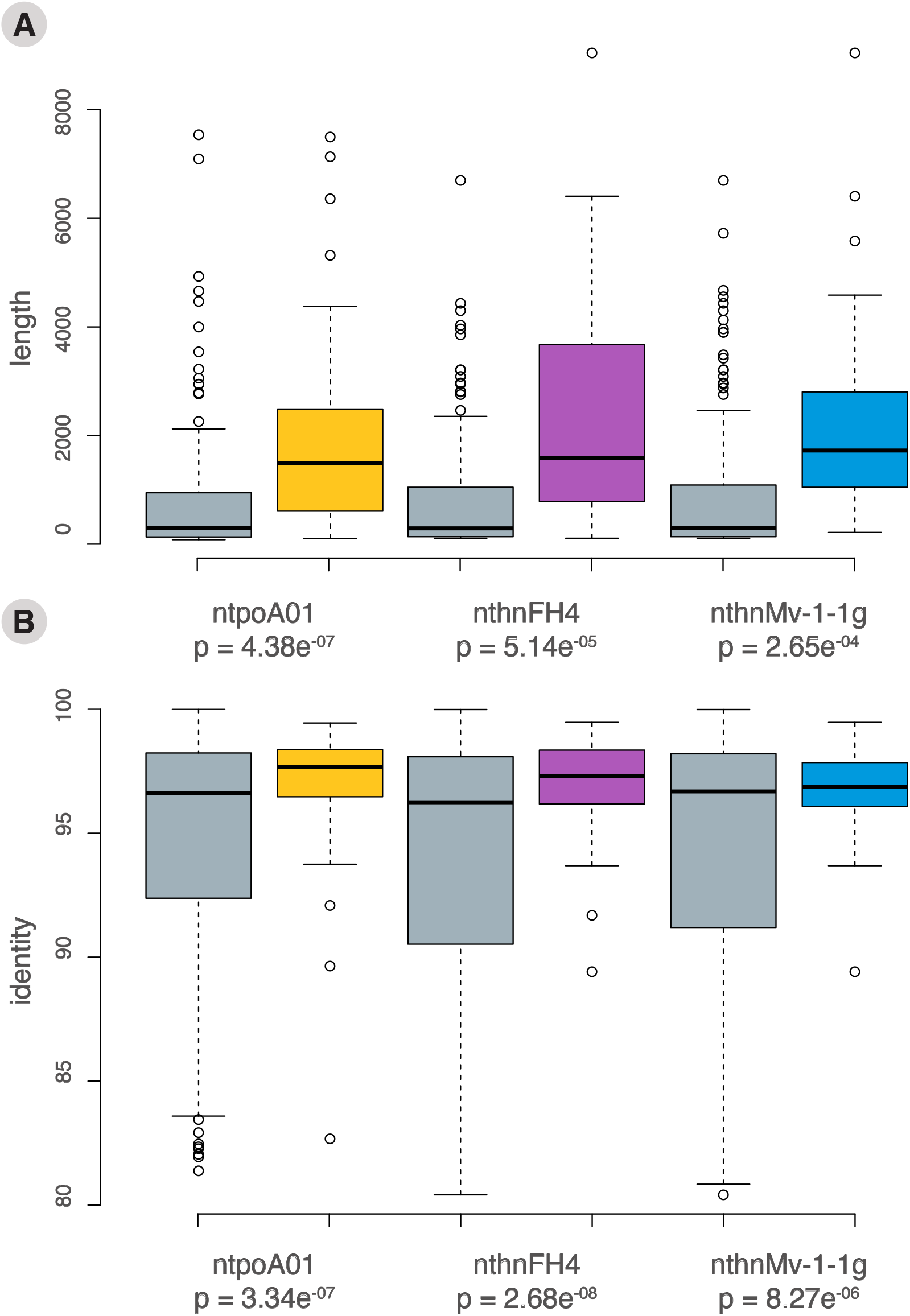
Differences in length (A) and sequence similarity (B, identity) between recombining homologous regions and non-recombining homologous regions greater than 100 bp. For both panels, the box plots (gray and colored for non-recombining and recombining homologous regions, respectively) show the mean of the distribution (horizontal bar), the 25 and 75% quartiles as a solid box, and the 5 and 95% quantiles as vertical lines. Below each comparison the *p*-value of the Student’s *t*-test is shown.

### Are there recombination hotspots?

In the three hybrid mitochondria analyzed a proportion of the homologous regions underwent multiple recombination events, while many others did not participate in non-allelic HR. As shown above, those homologous regions with greater length and sequence identity were more frequently involved in non-allelic HR leading to genomic rearrangements. Now, we wish to assess whether there are particular DNA tracts within those repeats that are more susceptible to recombination (*i*.*e*., recombination hotspots).

Given that two of the nthn somatic hybrids were constructed in independent experiments and their mtDNAs are chimeras of the same parental mitogenomes, we evaluated whether co-occurrence of recombination breakpoints within homologous regions was as expected by chance or not. For this, we performed a permutation test across all homologous regions present in the initial hybrid mitochondria meeting the necessary length and identity parameters that allows a HR event. The permutation test included a total of 207 homologous regions between *N. tabacum* and *H. niger* and showed that the observed co-occurrence of breakpoints in the same repeats was extremely unlikely (Fig. S8), suggesting an increased tendency to recombine at specific regions (Fig. 5).

**Figure 5.**
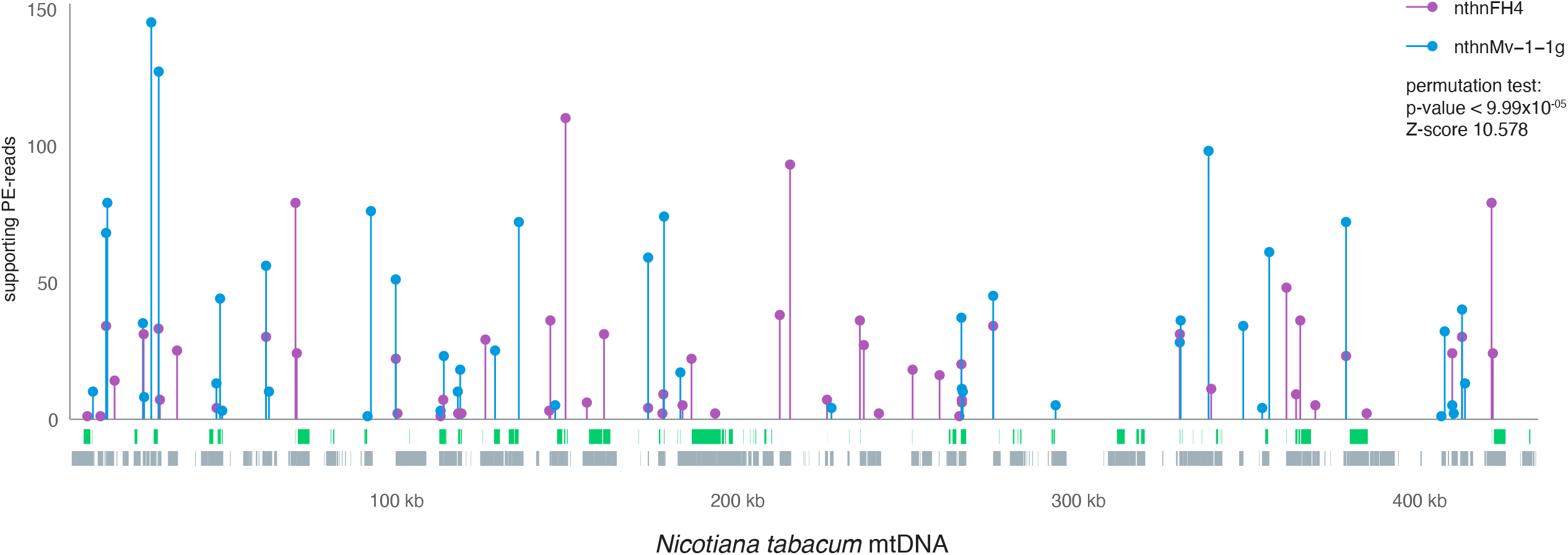
Recombination breakpoints in the *Nicotiana tabacum* mitogenome. Over the *X*-axis purple and blue lines depict the location of the recombination breakpoints over the tobacco genome for the nthnFH4 and nthnMv1-1-g somatic hybrids, respectively. Over the *Y*-axis, the frequency measured in supporting PE-reads is depicted. Below, the genes in green and the homologous regions between *N. tabacum* and *Hyoscyamus niger* mitogenomes in gray are shown.

## Discussion

Despite the early recognition of recombination as a factor in generating the structural diversity of plant mitogenomes (Palmer and Herbon, 1988; Lang *et al*., 1999), the replication and recombination processes in plant mitochondria are still not fully understood. Here, we focus on the recombination events that take place in the mitochondria of somatic hybrids after the protoplast fusion of sexually incompatible species. Findings in this experimental model might reflect the underlying recombination patterns in natural populations, which is difficult to study given that non-allelic recombination activity is generally low. Furthermore, this model might recapitulate the interaction between foreign and native mitogenomes upon the incorporation of horizontally transferred mitochondria from other plants, as proposed in the ‘fusion-compatibility model’ for mitochondrial HGT (Rice *et al*., 2013) light on the molecular mechanisms of foreign DNA incorporation into a native mitogenome. This study is also quite relevant to mitochondrial genome editing assays using nucleases, which entail DNA repair activities (Kazama *et al*., 2019; Arimura *et al*., 2020).

### Somatic hybrids are a useful model to study mitochondrial recombination in wild type plants

Somatic hybrid mitochondria carry hundreds of repetitive homologous regions, including numerous large imperfect repeats (Table S4, S6), offering a great opportunity to study multiple events and to evaluate the influence of repeat length and identity in non-allelic recombination. Wild type plant mitochondria generally employ homologous recombination pathways to repair DSBs in their DNA using as template an identical copy of the genome. If a DSB occurs within a repeated sequence, the non-allelic repeat is occasionally used as template, instead of the homologous repeat in another copy of the mitogenome. Only those HR events involving non-allelic repeated sequences can be detected by sequencing mitochondrial DNA and somatic hybrids present numerous products of recombination that can be precisely characterized.

Furthermore, it is possible to analyze a diverse population of mitogenomes from leaves and calli of a single somatic hybrid line that encompass a wide collection of chimeric molecules present in different cells. This is a great advantage of this approach because molecules with alternative configurations as a result of low frequency recombination events are often at low stoichiometries in plant mitochondria, even with less than one copy per cell and are prone to be lost during plant development (Arrieta-Montiel *et al*., 2001; Woloszynska and Trojanowski, 2009). This explains why the somatic hybrid ntpoA01 exhibits a higher number of interparental recombination events (117) than those observed in leaves of the cybrid plants nthn (∼50, Fig. 3).

### The assembly-free strategy is highly effective and sensitive to infer the number of recombination events

A known challenge when studying plant mitochondrial genomes relies on their assembly given the relatively large number of repeated sequences (Alverson *et al*., 2011; Gandini *et al*., 2019). This is particularly true in somatic hybrids because a large number of short and long homologous regions between the parental mitogenomes become repeats after mitochondria fusion. These repeats may produce complex assembly graphs, which hinder a correct characterization of mitochondrial genomes. Most times, particularly in those sequences assembled solely based on short-read sequencing, alternative conformations produced by recombination are ignored, and only the main conformation is presented in an assembled mitochondrial genome. To overcome this issue and inspired in previous works (Fritsch *et al*., 2014; Odahara, 2020), we developed a data analysis pipeline that does not require genome assembly. This strategy is powerful at recognizing multiple and independent recombination events that may occur across the same homologous region and it is also helpful at inferring recombination pathways.

The re-analysis of previously studied cybrids showed that the assembly-free approach is more sensitive as more recombination events were identified, in addition to the recognition of 95-98% of the events previously reported. Four of the five undetected events were missed because they are supported by two PE-reads of evidence, which suggests that our three PE-read threshold may be too restrictive. Given the high degree of constraints in the read mapping and filtering steps imposed by this pipeline, and that asymmetrical recombination frequently leads to the creation of subgenomic molecules at low stoichiometries, the existence of a single informative PE-read might be sufficient evidence for the identification of a true recombinant molecule. Considering a single chimeric read pair as evidence, 65 additional events can be inferred for the ntpoA01 mitogenome, 30 more for the nthnFH4, and 34 more for the nthnMv1-1g (Table S9), increasing to a greater extent the sensitivity of the assembly-free strategy.

### Mitochondrial recombination in somatic hybrids occurs mainly by BIR

By analyzing the products of interparental recombination, it is possible to infer the pathways that were involved. In most cases (65-87%), recombination was asymmetrical, with accumulation of only one of the two crossover products predicted by the dHJ pathway. These non-reciprocal recombination events are better explained by the BIR pathway (Fig. 2E) that results in a single product (Sakofsky *et al*., 2012; Sakofsky and Malkova, 2017). Reciprocal crossover products were observed in a low fraction of the non-allelic recombination events (0-6%) and may be the result of the dHJ pathway resolved with crossover. Alternatively, those reciprocal products could arise from two independent BIR events. Non-crossover products were identified in 3-21% of the HR events and may result from gene conversion via the SDSA and/or the dHJ (with no crossover) pathways. These products could also be explained by BIR with template switching from one repeat to the other during DNA synthesis (Sakofsky and Malkova, 2017).

The prevalence of the BIR pathway activity has been earlier recognized in asymmetrical events across short and intermediate repeats in plant mitochondria (Maréchal and Brisson, 2010; Sullivan *et al*., 2020). Studies on *Arabidopsis* RRR mutants, where factors such as WHY2 involved in HR were compromised, showed that asymmetrical recombination occurs frequently, leading to half-crossovers and mitochondrial rearrangements (Zaegel *et al*., 2006; Shedge *et al*., 2007; Cappadocia *et al*., 2010; Davila *et al*., 2011; Janicka *et al*., 2012; Miller-Messmer *et al*., 2012). These aberrant recombination events across intermediate repeats are presumably carried out through BIR and are suppressed by the nuclear machinery in wild type plants.

In contrast, it is widely accepted that long repeats undergo frequent reciprocal recombination in natural populations (Palmer and Shields, 1984; Arrieta-Montiel and Mackenzie, 2011). This type of recombination is compatible with the dHJ pathway leading to equimolar amounts of the crossover products (Chevigny *et al*., 2020). However, several studies found non-equimolar recombinational productsacross long repeats (Alverson *et al*., 2011; Wang *et al*., 2018; Dong *et al*., 2018; Sullivan *et al*., 2020). These observations could be explained either by a differential replication of the products, by asymmetrical recombination pathways that result in single products, such as BIR, or a combination of both processes. Likewise, our study shows that most recombination events after protoplast fusion lead to non-reciprocal products and involve long repeats with mean lengths of 1,998.53 bp, 2,074.34 bp, and 2,287.33 bp, for each of the three somatic hybrids, respectively. Thus, we provide strong evidence of the prevalence of asymmetrical recombination events across non-allelic repeats of all sizes, including long repeats, which are congruent with the BIR pathway.

Furthermore, the observations of rosette-like DNA structures, first by electron micrography, and lately in assembly graphs using long sequencing reads, suggest that BIR is involved in plant mitogenome replication (Backert and Börner, 2000; Gualberto and Newton, 2017; Jackman *et al*., 2020). This replication mechanism, *i*.*e*., recombination-dependent replication, has been well defined in the bacteriophage T4 and bacteria (Asai *et al*., 1994; Mosig, 1998), in which a ssDNA overhang invades a homologous site in another copy of the genome, priming DNA synthesis and eventually establishing a full replication fork (Maréchal and Brisson, 2010). The observed complex networks of plant organelle genomes can be explained by a prominent role of BIR in the recombination and replication processes (Manchekar *et al*., 2009; Maréchal and Brisson, 2010).

### BIR would also explain the bias deletions and the prevalence of a single allele in the hybrid mitogenomes

A notable feature from the studied somatic hybrids is the enormous amount of homologous parental sequences that were lost (∼40-50%) as opposed to almost complete retention of exclusive regions (78.10% to 99.76%). Parental content in cybrid mitogenomes was widely discussed in previous studies (Sanchez-Puerta *et al*., 2015; Garcia *et al*., 2019). The mitochondrial coding and non-coding sequences that were retained match the main parental component of the nuclear genome in the somatic hybrids, suggesting that the origin of the nuclear genome might impose restrictions on the number and origin of mitochondrial genes and their surrounding sequences. Taking into account these results, the authors presented two significant hypotheses: (1) adaptive forces could be eliminating duplicated sequences that may be unfavorable for mitochondrial and cellular performance; (2) a homologous recombination mechanism could be responsible for the neutral loss of an allele or parental form. The unexpected similarities in the biased sequence deletions in the three somatic hybrids analyzed reinforce the notion that duplicated regions tend to be eliminated by an asymmetrical HR pathway that favors the loss of a homologous region. As mentioned by García et al. (2019), this loss could be explained by a high activity of the BIR pathway after fusion of parental protoplasts.

### The co-occurrence of different recombination events suggests the existence of recombination hotspots

Our results strongly suggest the presence of recombination hotspots in plant mitochondria. In this study, we compared the location of the breakpoints in coincidental recombination events across non-allelic repeats in two somatic hybrids (nthn) and detected significant co-occurrence of HR events at the same breakpoints. However, the definition of recombination hotspot in plant mitochondria needs to be used with caution. Recombination hotspots in nuclear genomes are typically defined as those regions that suffer more DSBs and are repaired through recombination processes using the non-sister chromatid of homologous chromosomes in meiotic cells (Paul *et al*., 2016). In plant mitochondria, the multiple copies of the mitogenome are preferentially used as template for DSB repair and those recombination events are undetectable. Then, only those events between non-allelic repeats can be identified and the presence of hotspots can only be evaluated among the repeated sequences. In addition, ectopic recombination across short or intermediate repeats are inhibited in plant mitochondria, exhibiting low recombination rates, precluding the detection of hotspots in these regions. In conclusion, our ability to identify hotspots in plant mitochondria is restricted to large repeats (> 1 kb), which are frequently involved in non-allelic recombination events that can be observed. But bear in mind that the longer the repeat and the more similar in DNA sequence, the more likely to participate in non-allelic recombination, not because these repeats suffer more DSBs but because the nuclear factors do not inhibit so efficiently the recombination across regions with long tracts of homology and high identity. Thus, detection limitations and biases in non-allelic recombination events obscure the identification of regions that might suffer more DSBs in plant mitochondria, *i*.*e*., recombination hotspots. In line with this, earlier studies reported the existence of rearrangement hotspots in plant mitochondria as evidenced in Southern blot assays (Stern and Palmer, 1984; Kanazawa *et al*., 1998; Scotti *et al*., 2004; Zaegel *et al*., 2006). Those regions with increased rearrangement rates should not be confused with recombination hotspots. Those are likely large repeated regions prone to participate in non-allelic recombination events leading to rearranged molecules, as we observed in the three somatic hybrids (Fig. 4). In our analysis, we restricted our search of recombination hotspots to the location of the breakpoints among homologous regions susceptible to undergo non-allelic recombination. The detection of multiple independent recombination events occurring at the same breakpoints in two somatic hybrids indicates that some regions are more recombinogenic than others and can be considered recombination hotspots. Of course, there may be other hotspots across the mitogenomes outside of the repeats that cannot be detected with this approach.

### Non-homologous repair is rare in somatic hybrid mitogenomes

In this study, only 0-4 non-homologous repair events were identified in comparison to the 44-115 HR events detected in the three somatic hybrids analyzed. While in the nuclear genome of plants non-homologous end joining plays a leading role, non-homologous pathways are rarely observed in plant mitochondria (Devos *et al*., 2002; Chevigny *et al*., 2020). The latter could be a consequence of mitochondrial mechanisms to avoid error-prone recombination pathways inhibited by factors that favor HR pathways. Studies on Arabidopsis mutants in which HR pathways are inhibited uncovered the existence of microhomology-mediated repair (Arrieta-Montiel *et al*., 2009; Cappadocia *et al*., 2010; Janicka *et al*., 2012; Zampini *et al*., 2016). These non-homologous pathways were involved in rare mitogenome rearrangements (Hunt and Newton, 1991), the truncation of mitochondrial genes (Kadowaki *et al*., 1990; Newton *et al*., 1990), and in the reversion of mutant phenotypes (Feng *et al*., 2009).

### Somatic hybrids as a model to study sequence acquisition after HGT events

Plant cells contain mitochondria with an uneven distribution of the mitochondrial DNA. Some mitochondria possess complete genomes, some harbor subgenomic molecules, and others are devoid of genetic material (Sheahan *et al*., 2005). After protoplast fusion and before starting cell division, the mitochondria undergo massive mitochondrial fusion (MMF), mixing the population of mitochondrial genomes in approximately 24 hours. MMF was first observed in somatic hybrids and is believed to act as a control mechanism, providing the opportunity for all sub-genomes to interact recombining and repairing DNA before cell replication (Sheahan *et al*., 2005; Seguí-Simarro *et al*., 2008; Paszkiewicz *et al*., 2017). In that sense, somatic hybrids can mimic the process of HGT described by the fusion-compatibility model (Rice *et al*., 2013) in which (1) foreign mitochondria enter a recipient cell, fuse with native mitochondria, and their genomes recombine; (2) during the subsequent cell divisions, foreign sequences should be replicated, along with the native genome; (3) the transformed cells must become part of the germline and be inherited by the next generation. The high frequency of BIR observed in the somatic hybrids remarks that foreign sequences are likely integrated and replicated through this recombination pathway. Homologous regions between the foreign and the host mitogenomes will undergo asymmetrical recombination, resulting in the loss of one of the homologous regions, either randomly or shaped by natural selection based on nuclear-cytoplasmic compatibilities. In contrast, those mitochondrial sequences with little or no similarity will remain as long as they continue to be replicated and perpetuated under the whims of genetic drift.

## Abbreviations

(BIR): break-induced replication
(CO): crossover
(DSBs): DNA double-strand breaks
(RRR): DNA recombination, repair, and replication
(dHJ): double Holliday junction
(HR): homologous recombination
(MMF): massive mitochondrial fusion
(nthn): *Nicotiana tabacum* and *Hyoscyamus niger* somatic hybrid
(ntpo): *Nicotiana tabacum* and *Physochlaina orientalis* somatic hybrid
(non-CO): non-crossover
(PE): paired-end
(SDSA): synthesis-dependent strand annealing

## Supplementary data

The following supplementary data are available at JXB online.

**Table S1**. Total reads and datasets created by differential read filtering.

**Table S2**. Comparison of gene content in the mitochondrial genomes *of Nicotiana tabacum* (NC_006581) and *Physochlaina orientalis* (NC_044153) and their contribution to the ntpoA01 somatic hybrid.

**Table S3**. Summary of parental sequence contribution and recombination events in the somatic hybrids ntpoA01, nthnFH4, and nthnMv-1-1g.

**Table S4**. Homologous regions between the mitogenomes of *Nicotiana tabacum* (nt) and *Physochlaina orientalis* (po).

**Table S5**. Recombination events in the ntpoA01 somatic hybrid mitochondria.

**Table S6**. Homologous regions between the mitogenomes of *Nicotiana tabacum* (nt) and *Hyoscyamus niger* (hn).

**Table S7**. Recombination events in the nthnFH4 somatic hybrid mitochondria.

**Table S8**. Homologous recombination events in the nthnMv-1-1g somatic hybrid mitochondria.

**Table S9**. Recombination events supported by less than 2 reads.

**Figure S1**. Southern blot experiments using the mitochondrial genes *atp1, cob, cox1*-exon 2 or *cox1*-intron as probes to test the chimeric status of the mitochondria in somatic hybrids between tobacco (Nt) and *Physochalina orientalis* (Po).

**Figure S2**. Schematic illustration describing the methodology to collect recombination evidence and to infer regions contributed by each parent in somatic hybrid mitochondria.

**Figure S3**. Patterns of PE read alignment to infer recombination pathways in somatic hybrid mitochondria.

**Figure S4**. Graphical evidence for each recombination event identified in the ntpoA01 mitogenome.

**Figure S5**. Graphical evidence for each recombination event identified in the nthnFH4 mitogenome.

**Figure S6**. Graphical evidence for each recombination event identified in the nthnMv-1-1g mitogenome.

**Figure S7**. Differences in length and sequence similarity between recombining and all the non-recombining homologous regions.

**Figure S8**. Permutation test of a total of 207 homologous regions between *N. tabacum* and *H. niger* and the observed co-occurrence of recombination across the same repeats setting tobacco or *H*.*niger* as the reference.

**Methods S1**. Somatic hybrid production and Southern blot experiments.

**Methods S2**. Inferring recombination pathways.

**Data S1**. The plastid genome of the somatic hybrid ntpoA01.

## Author Contributions

CLG contributed with conceptualization, design methodology, software, formal analysis, investigation, interpretation, and data curation, drafting, writing and revising of the work, and with the visualization process. LEG contributed with the interpretation of data and with the review and editing of writing. CCA, LFC, SK and DG all contributed to the creation and maintenance of the somatic hybrids (methodology) and with the manuscript revising. MVSP contributed with the conceptualization, the supervision, the review, editing and final approval of writing, the project administration, and the funding acquisition.

## Conflict of interest

No conflict of interest declared.

## Funding Statement

This research received support from grant PICT2017-0691 from FONCyT to M.V.S.P.

## Data Availability

The data that support the findings of this study is openly available in the SRA archive at https://www.ncbi.nlm.nih.gov/sra, under the project PRJNA899353.

## Notes

### Competing Interest Statement

The authors have declared no competing interest.

